# National Population and Cost Implications of Treatment with Icosapentyl Ethyl in the United States: An Assessment Based on the REDUCE-IT trial

**DOI:** 10.1101/466649

**Authors:** Rohan Khera, Javier Valero-Elizondo, Anshul Saxena, Salim S Virani, Harlan M Krumholz, Khurram Nasir

## Abstract

Icosapent ethyl, an omega-3-fatty acid, was associated with improved cardiovascular outcomes in individuals with elevated levels of serum triglycerides and at a high-risk of adverse cardiovascular events in the recently completed REDUCE-IT trial. We applied the eligibility criteria of the REDUCE-IT trial to a nationally representative sample of individuals in the United States captured in the National Health and Nutrition Examination Survey (NHANES) for a 6-year period (2009-2014) and estimated the number of individuals nationally that would be potentially eligible for treatment with icosapentyl ethyl. We found that nearly 3 million US adults would potentially be candidates for this therapy. Further, based on the list cost of the drug, if all eligible individuals are treated, the additional cost to the US healthcare system will be almost $9 billion a year.

## BACKGROUND

**I**n the REDUCE-IT (Reduction of Cardiovascular Events Outcomes) trial – a multicenter, randomized, placebo controlled trial – treatment with icosapent ethyl, an omega-3-fatty acid among individuals with high cardiovascular risk and elevated triglycerides in the setting of concurrent statin use was associated with 25% reduction in major adverse cardiovascular events, including cardiovascular death, nonfatal myocardial infarction, stroke, revascularization and unstable angina requiring hospitalization.^1,2^ The study follows other evidence of effective lowering of serum levels of triglycerides with the use of this agent, in addition to improvement in atherogenic lipid profile.^3^ Given the large unmet need for prevention of adverse cardiovascular outcomes among those with established cardiovascular disease or at high risk for developing the disease, it is critical to evaluate the implications of REDUCE-IT for the national US population.

## METHODS

In the National Health and Nutrition Examination Survey (NHANES) for 2009 through 2014 – a large population based sample of non-institutionalized individuals from the United States – we identified all individuals that satisfied the study selection criteria for the REDUCE-IT trial. Briefly, we identified individuals ≥45 years of age with a history of atherosclerotic cardiovascular disease (ASCVD) (group 1) and those ≥50 years without ASCVD but with diabetes mellitus with one additional cardiovascular risk factors, including hypertension, reduced high density lipoproteins, cigarette smoking, advanced age, and renal dysfunction (group 2).^2^ For each group, those with a triglyceride ≥150, but <500 milligrams per deciliter (mg/dL), LDL of ≥ 40 to < 100 mg/dL, and ongoing statin therapy were further selected, and those with major exclusions were excluded. The major study exclusions are listed in **Figure 1**.

**Figure 1.**
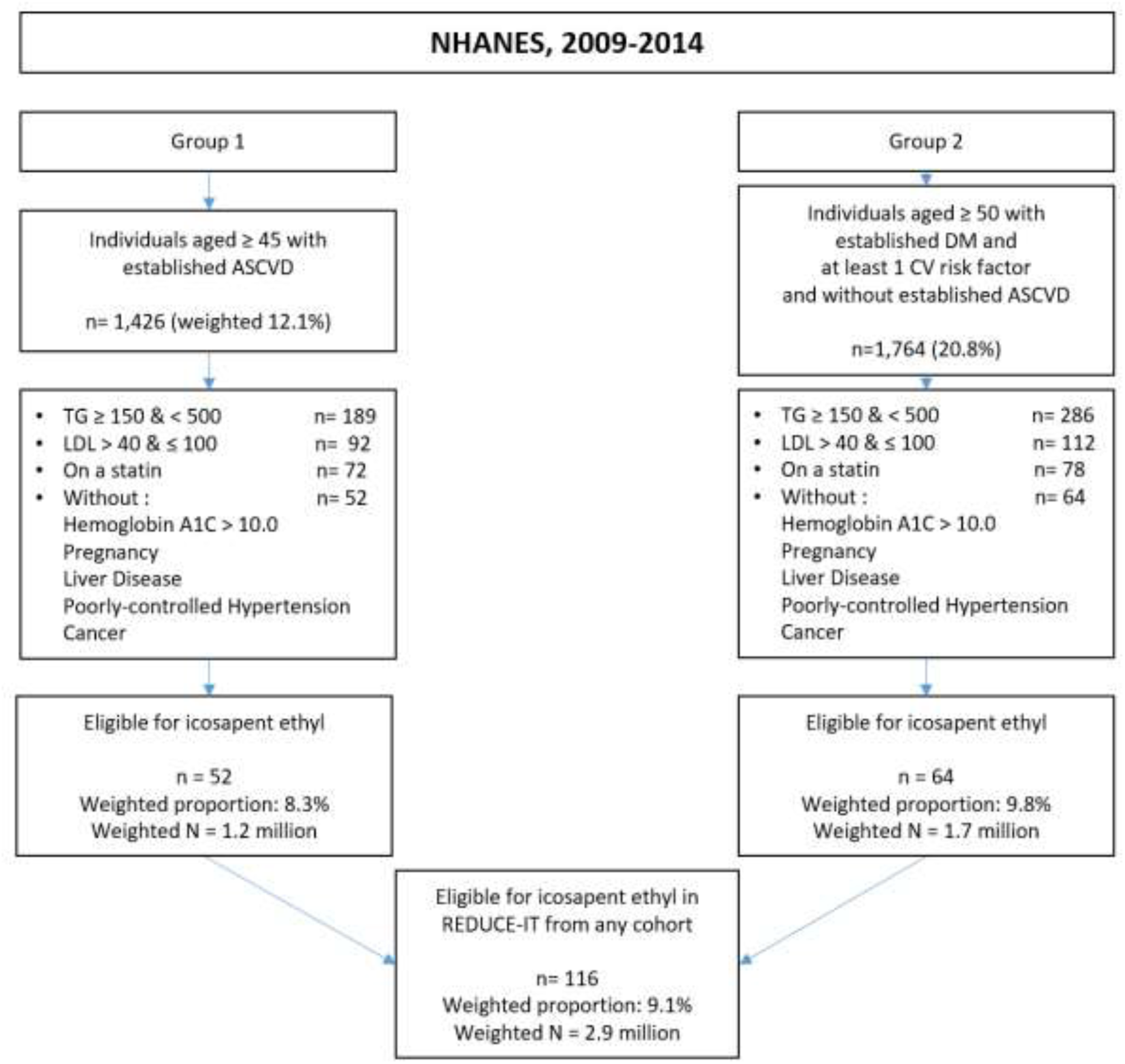
Selection of individuals eligible for treatment with icosapentyl ethyl.

We used survey-specific analyses to obtain nationally-representative weighted estimates. These analyses specifically accounted for the complex sampling design of the NHANES, including the stratification and clustering of data. We combined 3 cycles of the NHANES – 2009-2010, 2011-2012, and 2013-2014 – to obtain more reliable estimates. The survey weights for the three cycles were divided by 3 to obtain modified survey weights such that the weighted estimates represented national estimates during an average year during this period. Estimates for the cost of treatment of all individuals eligible for icosapentyl ethyl were based on its retail cost of $242.32 per month on November 8, 2018 for a daily 4 gram dose.^4^ Estimated cost of therapy for treating all eligible individuals nationally and the 95% confidence intervals for this estimate were calculated by multiplying the monthly list cost with the estimate for the total number of individuals eligible for therapy and the corresponding 95% confidence intervals for the eligible population estimate.

## RESULTS

In NHANES 2009-2014, 1426 individuals, representing 12.1% of the adult population over age 45 years had ASCVD, and 1764 individuals, representing 20.8% of the adult population over 50 years, did not have ASCVD, but had diabetes and one or more cardiovascular risk factors. Among these, 52 or 8.3%, representing 1.2 million (95% CI 0.8 million – 1.9 million) nationally would qualify for icosapent ethyl based on ASCVD and 64 or 9.8%, representing 1.7 million individuals nationally (95% CI 1.3 million – 2.3 million) would qualify based on diabetes and risk factors. Collectively, 116 individuals in the sample, representing 2.9 million (2.2 million – 3.8 million) individuals nationally would qualify for therapy with icosapent ethyl (**Figure 1**).

Among those eligible for therapy, mean age was 54 years (95% CI, 39 – 68 years) and an equal proportion were men and women. A majority of these patients had private health insurance (63.1%), while a third were publicly insured (**Table 1**). Notably, an additional 839,515 individuals (95% CI, 0.6 million – 1.2 million) had lipid levels in therapeutic range for treatment but require initiation of a statin before further consideration for treatment with icosapentyl ethyl.

**Table 1.**
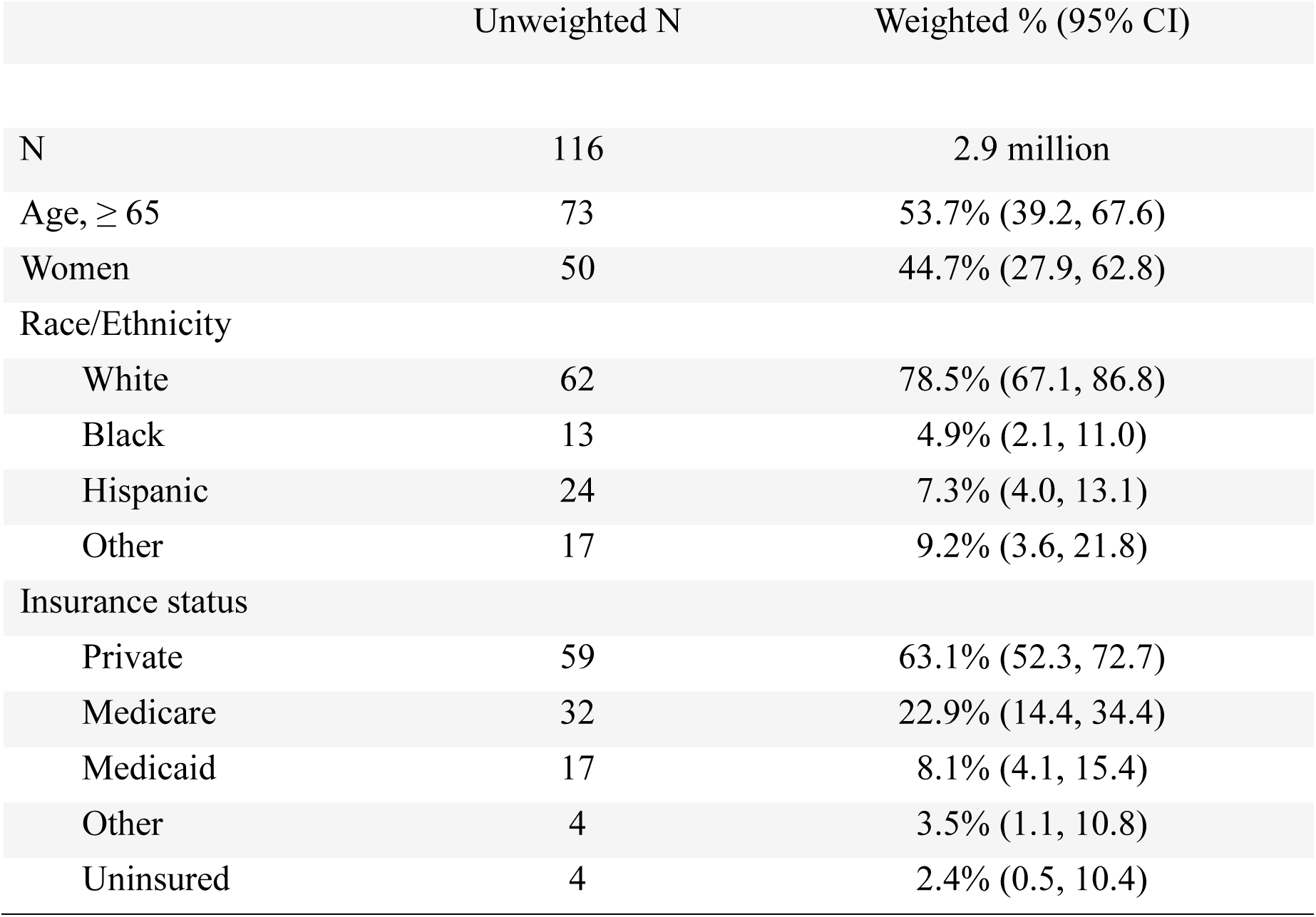
Characteristics of individuals qualifying for therapy with icosapentyl ethyl.

The treatment of all eligible individuals would represent an overall cost of $709 million per month (538 million – 928 million) assuming a monthly cost of $242.32 for a 4g dose of icosapentyl ethyl,^4^ the dose evaluated in the REDUCE-IT trial.^2^ The treatment of just those with ASCVD would cost $291 million per month ($183 – 455 million), while treatment of those at risk for ASCVD would cost $418 million per month ($316 – 548 million).

## DISCUSSION

Nearly 3 million US adults with or at high risk for ASCVD have elevated serum lipid levels despite treatment with a statin and would qualify for treatment with icosapentyl ethyl to lower their risk of adverse cardiovascular events. Moreover, nearly a million additional individuals who currently are not being treated with statins but would meet eligibility criteria for treatment with icosapentyl ethyl based on their current lipid profile. Extending this therapeutic option to all eligible adults would result in $709 million in healthcare spending every month, or a total of $8.5 billion annually.

Our assessment should be interpreted in the light of a few limitations. The overall number of individuals captured in NHANES is small. However, our estimates are reliable and comply with the suggested relative standard error of less than 30% suggested for analyses with NHANES. Moreover, our sample size is comparable to similar assessments in previous studies. The measurement of high-sensitive C-reactive protein was performed only for 2 years of our 6-year study period, but the addition of C-reactive protein thresholds to our patient selection algorithm did not identify any additional individuals in these 2 years. However, we may potentially underestimate the eligibility of US adults if some individuals in the latter years of the study had elevated high-sensitivity C-reactive proteins but did not meet additional criteria for inclusion. Finally, costs of therapy are based on the list price of the drug and do not account for potential rebates that are likely to be available with the widespread utilization of the drug.

In conclusion, almost 3 million US adults would be candidates for therapy with icosapentyl ethyl and if all are treated the additional cost to the US healthcare system will be almost $9 billion a year if list costs are maintained.

## REFERENCES

1. Amarin corporation. REDUCE-IT^TM^ cardiovascular outcomes study of vascepa^®^ (icosapent ethyl) capsules met primary endpoint. Available at: https://investor.amarincorp.com/news-releases/news-release-details/reduce-ittm-cardiovascular-outcomes-study-vascepar-icosapent. Accessed November 4, 2018.

2. Bhatt DL, Steg PG, Brinton EA, et al. Rationale and design of REDUCE-IT: Reduction of Cardiovascular Events with Icosapent Ethyl-Intervention Trial. Clin Cardiol. 2017;40(3):138-148.

3. Mosca L, Ballantyne CM, Bays HE, et al. Usefulness of Icosapent Ethyl (Eicosapentaenoic Acid Ethyl Ester) in Women to Lower Triglyceride Levels (Results from the MARINE and ANCHOR Trials). Am J Cardiol. 2017;119(3):397-403.

4. GoodRx. Vascepa drug prices. Available at: https://www.goodrx.com/vascepa. Accessed, November 3, 2018.

